# Aberrant Hippo-YAP/TEAD signaling drives malignant transcriptional reprogramming in external auditory canal squamous cell carcinoma

**DOI:** 10.1101/2025.08.27.672711

**Authors:** Kuniaki Sato, Noritaka Komune, Mayumi Ono, Takahiro Hongo, Takafumi Nakano, Kensuke Koike, Shinsaku Itoyama, Kenichi Taguchi, Koshi Mimori, J. Silvio Gutkind, Muneyuki Masuda, Takashi Nakagawa

## Abstract

**Purpose:** External auditory canal squamous cell carcinoma (EACSCC) is an extremely rare malignancy. The molecular characteristics and evidence-based therapeutic strategies of EACSCC still remain to be elucidated.

**Experimental Design:** Comprehensive analyses of RNA sequencing (RNA-seq) and ChIP sequencing (ChIP-seq) utilizing YAP and H3K27Ac antibodies were performed in primary EACSCC and noncancerous ear skin samples. Functional experiments were performed in EACSCC-derived cells and Head and Neck Squamous Cell Carcinoma (HNSCC) cells *in vitro* and *in vivo*. Immunohistochemical staining of primary EACSCC tissues as well as survival analysis were conducted.

**Results:** RNA-seq indicated hyperactivation of YAP/TEAD-mediated transcriptional programs in EACSCC. H3K27Ac ChIP-seq suggested gained accessibility for transcription factor (TF) binding sites for TEAD, AP-1 and PITX TFs in EACSCC, and presence of EACSCC-specific super enhancers (SEs). YAP-bound SEs were involved in oncogenic transcription, including EGFR signaling. Small molecule TEAD inhibitor (smTEADi) VT104 showed significant suppression of proliferation and clonogenicity in EACSCC cells. Importantly, smTEADi not only inhibited YAP-TEAD interaction but also induced YAP-PITX2 binding, suggesting that PITX2 could represent an alternative partner TF of YAP under TEAD-inhibited conditions. Knockdown of PITX2 inhibited cell growth and migration of EACSCC and HNSCC cells, whereas overexpression of PITX2 induced expression of cell cycle, stemness, and EMT genes, as well as YAP/TAZ-TEAD target genes, and promoted tumor growth *in vivo*. Nuclear YAP and PITX2 expression were significantly correlated with poor prognosis of EACSCC patients.

**Conclusions:** This study highlighted the hyperactivation of the YAP-TEAD/PITX2 transcriptional program and its potential as a therapeutic target in EACSCC.

**Translational Relevance:** External auditory canal squamous cell carcinoma (EACSCC) is an extremely rare malignancy related to chronic tissue damage and inflammation. Due to its rarity, the molecular characteristics of EACSCC are poorly understood, and evidence-based therapeutic strategies are not fully developed. Here, we provide evidence of hyperactivation of YAP/TEAD-driven transcriptional programs in EACSCC, utilizing comprehensive analyses of RNA-seq and YAP/H3K27Ac ChIP-seq in clinical tissue samples, as well as *in vitro* and *in vivo* experiments. In addition, our data suggest that the PITX2 transcription factor (TF) could represent an alternative partner TF of YAP under TEAD-inhibited conditions, which may rescue oncogenic transcription of TEAD. Importantly, YAP and PITX2 are co-expressed in EACSCC and predict poor prognosis of EACSCC patients. Our results provide a rationale for YAP-hyperactivation in EACSCC and contribute to a better understanding of this malignancy and the development of new therapeutic strategies.

## Introduction

The notion “*Tumors are wounds that do not heal*” suggests the close links between cancer, inflammation, tissue damage, and wound healing (1). The similarity between cancer onset/progression and wound healing process has long been described, in which signaling pathways and microenvironmental changes are shared in many aspects (2-4). Importantly, recent studies revealed that non-genetic alterations, or “epigenetic reprogramming,” including chromatin remodeling and aberrant transcription, may regulate these processes (5,6). One of the most important molecules for this reprogramming is Yes-associated protein (YAP), a transcriptional co-activator negatively regulated by the Hippo pathway (7). Mechanistically, YAP drives transcription of target genes cooperatively with partner transcription factors (TFs), especially with TEA domain family members (TEAD), in response to mechanical and microenvironmental stress (8,9). Indeed, several lines of evidence have shown crucial roles of YAP in wound healing and tissue regeneration (10-12) as well as carcinogenesis (13-15), suggesting that YAP acts as a transcriptional driver of malignant transformation and tumor progression in multiple tumor types.

External auditory canal squamous cell carcinoma (EACSCC) is an extremely rare malignancy, which is reportedly diagnosed in one to six out of a million individuals annually, arising from the squamous epithelium of the external auditory canal (16). The biological and molecular characteristics of EACSCC, as well as evidence-based therapeutic strategies for this malignancy, have not yet been fully established due to its rarity. Unlike head and neck squamous cell carcinoma (HNSCC) or skin basal cell carcinoma, smoking or UV exposure is not evident in the mutational signature of EACSCC, as we previously reported by whole exome sequencing (17), and the presence of high-risk human papillomavirus (HPV) infection is extremely rare (18). Clinically, risk factors of EACSCC include chronic tissue damage and inflammation induced by mechanical stimuli (e.g., habitual ear-picking especially in Eastern Asian countries) (19), which may suggest that continuous and aberrant hyperactivation of YAP-driven regenerative transcriptional programs could drive malignant initiation and progression of EACSCC. Moreover, we previously showed that mutations recurrently observed in EACSCC included loss-of-function (LOF) mutations of FAT atypical cadherin 1 (*FAT1*), which is an upstream regulator of the Hippo pathway (17). LOF of FAT1 results in Hippo pathway dysfunction and subsequent YAP hyperactivation, as we previously reported in HNSCC (20). Taken together, we hypothesized that YAP-driven transcriptional programs could be a crucial oncogenic driver of EACSCC.

Here, we report the transcriptomic and epigenetic aberrations of EACSCC. Comprehensive analysis of RNA sequencing (RNA-seq) and chromatin immunoprecipitation sequencing (ChIP-seq) in clinical EACSCC samples and noncancerous ear skin tissues uncovered hyperactivation of YAP and downstream pathway aberrations. Additionally, we revealed a potential alternative transcriptional machinery of YAP under TEAD-inhibited conditions, and utilizing multiple experimental strategies, including small molecule TEAD inhibitor and EACSCC-derived cells, we investigated the clinical significance of YAP-hyperactivation in EACSCC.

## Materials and Methods

### Ethics Statement

The protocol of this study was reviewed and approved by the institutional review boards and ethics committees of Kyushu University (Protocol Number: 700-2 and 30-268). All experiments with human samples were conducted according to the principles expressed in the Declaration of Helsinki.

### Patients and sample collection

All patients diagnosed as EACSCC and treated at Kyushu University Hospital Department of Otolaryngology from September 2015 to March 2019, were provided written informed consent and enrolled in this study. Primary tumor samples were collected by tissue biopsy or surgical resection. Noncancerous ear skin tissues were collected from surgically resected tissues.

### RNA extraction, reverse transcription, and qPCR

Total RNA was isolated from EACSCC and noncancerous ear skin tissues using a NucleoSpin RNA Plus kit (Macherey-Nagel, Germany). Reverse-transcription was performed using a PrimeScript™ RT Master Mix (Takara Bio, Japan) followed by qPCR was performed in a CFX96 system (Bio-Rad Laboratories, USA) with TB Green® Premix Ex Taq™ II (Tli RNase H Plus) (Takara Bio). The primers used in this study are listed in Table S2. β-actin was used as an internal control. The mRNA expression levels were calculated using the 2-ΔΔCt method.

### RNA sequencing and analysis

RNA extracted from EACSCC and skin tissues was sequenced on a DNBSEQ-G400 sequencer at Beijing Genomics Institute (Shenzhen, China). The sequencing reads were aligned to the human reference GRCh38/hg38 genome by STAR v2.7.9 using Gencode v38 annotations. Gene count tables were generated using RSEM. Downstream analyses were carried out using R v4.4.1 (The R Foundation for Statistical Computing, Austria). Normalization of the read count data and the detection of differentially expressed genes (DEGs) between EACSCC and skin were carried out with DESeq2 v1.10.1. For sample clustering and principal component analysis, genes with zero counts across all samples were removed from the analysis. Gene Ontology (GO) analysis of DEGs was performed using Molecular Signature Database (MSigDB) Hallmark pathways, GO Biological Process and KEGG pathways in the R package enrichR. Gene set enrichment analysis (GSEA) was performed using GSEA v4.3.2 (21). Single-sample GSEA (ssGSEA) was performed in GenePattern (www.genepattern.org). TF activity prediction was performed using the R package decoupleR v2.9.1 (22). YAP/TAZ-TEAD transcriptional target signature gene sets were obtained from the previous studies (23,24).

### ChIP sequencing and analysis

Frozen tumor samples and skin tissue samples were sent to Active Motif (Carlsbad, CA, USA) for ChIP reaction, library preparation, and sequencing. Crosslinked cells were solubilized using lysis protocol, sonicated, and immunoprecipitated with antibodies for histone 3 lysine 27 acetylation (H3K27Ac) (#39133, Active Motif) and YAP (#Y1200-01D, US Biological). Immunoprecipitated chromatin samples were sequenced on an Illumina NextSeq 500 sequencer. The 75bp single-end raw reads were aligned to the human reference genome GRCh38/hg38 using BWA v0.7.12. Only reads aligned with no more than 2 mismatches and mapped uniquely to the genome are used in the subsequent analysis. PCR-duplicated reads were removed. Peaks were identified using MACS2 v2.1.0 (p-value cutoff = 1e-7), and respective input samples were used as background. For visualization of called peaks and regions, deeptools and pyGenomeTracks were used. The ROSE algorithm was applied for the detection of SEs using default parameters (25). Calculation of overlapping between SE regions was performed using bedtools *multiinter*, and the regions with lengths less than 3kb were filtered out. SE regions observed in at least two EACSCC samples, but never observed in noncancerous skin samples, were defined as EACSCC-specific SEs. Enhancer-to-gene interactions in the detected enhancers and SEs were estimated using GeneHancer v5.24, a comprehensive enhancer/promoter database of the human genome (26). Regions annotated as promoters were removed. Nucleosome-free regions (NFRs) in H3K27Ac peak regions were estimated using the HisTrader algorithm as previously described (27). Detected YAP peaks were annotated to putative transcriptional target genes using GREAT (28) with *Basal plus Extension* option (2kb upstream and 2kb downstream, distal up to 1,000kb options). Transcription factor binding motif discovery analysis in 200bp regions around YAP peak summits and motif enrichment analysis in NFRs were performed using HOMER v5.1 (29) with the default database included in HOMER and HOCOMOCO v13 database (30).

### Cell lines and culture

Human HNSCC cell lines SCC9 and FaDu were obtained from the American Type Culture Collection, and HSC4 was obtained from the Japanese Collection of Research Bioresources. SCEACono2 was established from a primary EACSCC as previously described (31,32). Cells were maintained in Dulbecco’s modified Eagle’s medium (DMEM)/F12 (Sigma Aldrich, USA), supplemented with 10% fetal bovine serum (FBS: Sigma Aldrich), 0.1 mM MEM nonessential amino acids solution (Thermo Fisher Scientific, USA), 1 mM sodium pyruvate (Thermo Fisher Scientific), 2 mM L-glutamine (Thermo Fisher Scientific), and 1% antibiotic antimycotic mixed solution (Nacalai Tesque, Japan) at 37°C with 5% CO_2_.

### Western blotting

Cultured cells were lysed with RIPA buffer (Thermo Fisher Scientific) containing protease inhibitor (Sigma Aldrich) on ice. Protein concentration was determined using a BCA protein assay Kit (Takara Bio). A total of 30 *µ*g protein/lane were loaded on 10% Mini-PROTEAN TGX Tris-Glycine gels (Bio-Rad Laboratories), separated by SDS-PAGE, and then transferred to PVDF membranes. After blocking with 5% non-fat milk in TBS-T, the membranes were incubated overnight at 4°C with the following primary antibodies; PITX2 (1:500, #PA5-98817, Thermo Fisher Scientific), YAP (1:1000, #14074, Cell Signaling Technology, USA), TEAD1 (1:1000, #12292, Cell Signaling Technology, USA), CD44 (1:1000, #3570, Cell Signaling Technology), CDH2 (1:1000, #13116, Cell Signaling Technology), Snail (1:1000, #3879, Cell Signaling Technology) and β-actin (1:1000, #4967, Cell Signaling Technology). The membranes were then incubated with the goat anti-rabbit IgG/horseradish peroxidase-conjugated secondary antibody (1:2000, #7074, Cell Signaling Technology) for 6 1 h at room temperature. For the detection of protein bands, the Clarity ECL Western Blotting Substrates and ChemiDoc MP Imaging system (Bio-Rad Laboratories) were used.

### Co-immunoprecipitation (Co-IP)

Co-IP was performed using SureBeads Protein A/G magnetic beads (Bio-Rad Laboratories) according to manufacturer’s protocols. Briefly, cells at 70% confluency were lysed with Pierce IP Lysis Buffer (Thermo Fisher Scientific) on ice. The total cell lysate was incubated with YAP antibody (1:100, #14074, Cell Signaling Technology) or IgG for 2 h at room temperature, followed by incubation with magnetic beads at 4°C overnight. The immunoprecipitants were washed with PBS containing 0.1% Tween 20 (PBS-T) three times. Proteins bound to the beads were eluted using 1x SDS sample buffer (Bio-Rad Laboratories) at 98°C for 5 min. Eluted samples were subjected to SDS-PAGE and western blotting with input samples.

### Plasmid transfection and generation of stable expressing cells

pCMV6 control plasmid and PITX2 overexpressing plasmid (#RC204179) were purchased from Origene. Transfection was performed using PEI Max (Polyscience, USA) according to the manufacturer’s instructions. 24h after transfection, the cells were subjected to antibiotic selection in medium containing G418 (Thermo Fisher Scientific). Cells stably expressing the recombinant protein were pooled for experiments.

### siRNA-mediated knockdown experiments

Cells were seeded onto 6-well plates (2×10^5^ cells/well) and incubated overnight. 20nM pooled siRNAs for PITX2 (#sc44016, Santa Cruz Biotechnology, USA) and negative control siRNA (#sc37007, Santa Cruz Biotechnology) were transfected to cells using Lipofectamine RNAiMAX (Thermo Fisher Scientific) according to the manufacturer’s protocol. After 48h of transfection, the cells were harvested for RNA/protein extraction. Transfected cells were used for downstream analyses.

### Cell proliferation assay

Cell viability was detected using a Cell Counting Kit-8 (CCK-8) cell proliferation assay (Dojindo Molecular Technologies, Inc., Japan) according to manufacturer’s protocol. Briefly, the cells were seeded onto 96-well plates (2×10^3^ cells/well) and cultured overnight at 37°C. 10 *µ*l CCK-8 solution was added at 24, 48, 72 and 96 h timepoints after seeding. The absorbance at 450 nm was measured.

### Migration assay

Cells were treated with PITX2 siRNA or control siRNA for 24 h, then 2×10^5^ cells/well were seeded onto an upper chamber of 24-well Transwell insert (Corning, USA) and incubated for 18 h. The cells that migrated into the lower side of the filter were fixed and stained with DAPI (Vector Laboratories, Inc., USA) and then counted under a fluorescence microscope.

### Scratch wound healing assay

Cells were seeded onto 6-well plates at a density of 2×10^5^ cells/well and grown to confluence, then transfected with PITX2 siRNA or control siRNA overnight. Scratches were made with a 1ml plastic pipette tip across the diameter of each well, and cells were cultured in serum-free conditions. After 24h, the cell-migrated areas in the scratch were quantified using ImageJ.

### Sphere formation assay

Cells were seeded onto 6-well ultra-low attachment plates (Corning) at a density of 5,000 cells/well in Ham’s F12 containing 20 ng/ml EGF (#AF-100-15, PeproTech, USA), 20 ng/ml bFGF (#100-18B, PeproTech), and B27 Supplement (#17504044, Invitrogen). Triplicate wells were prepared for each cell. After 2 weeks, visible spheres were manually counted using a microscope.

### Clonogenic assay

Cells were plated onto 24-well plates at a density of 2,500 cells/well in triplicate and cultured at 37ºC under 5 % CO2 overnight. Subsequently, cells were treated with VT104 TEAD inhibitor (#HY-134956, MedChemExpress, USA) at 10-10,000 nM concentrations or DMSO for 7 days and then fixed with 10% acetic acid in 100% methanol for 15 min. Fixed cells were stained with 0.5% Crystal Violet in 20% methanol for 1 h. Stained colonies were washed with distilled water and dried, then scanned using a document scanner. For quantification, Crystal Violet was dissolved in 10% acetic acid in water, and absorbance at 595nm was measured in a plate reader.

### Histological analyses and Immunohistochemical staining

The protein expression levels of YAP1 and PITX2 in the primary EACSCC tissues were evaluated by immunohistochemistry (IHC) as previously described (14). The antibodies used in this analysis were as follows: YAP (#WH0010413M1, Sigma Aldrich), PITX2 (#PA5-98817, Thermo Fisher Scientific). The histology and the results of IHC were independently reviewed by three experienced pathologists (T.H., T. Nakano and T.K.). For IHC staining of mouse xenograft tumor tissues, formalin-fixed, paraffin-embedded specimens obtained from tumors were stained with PITX2 and Ki 67(#M7240, Dako, USA) as previously described (31).

### Murine xenograft models

All animal procedures were performed in compliance with the Guidelines for the Care and Use of Experimental Animals established by the Committee for Animal Experimentation of Kyushu University. Five-week-old female BALB/c nu/nu mice were purchased from Japan SLC and maintained under specific pathogen-free conditions. 1×10^6^ SCEACono2 cells expressing PITX2 or control cells were subcutaneously implanted. Tumor sizes were measured every three days using a Vernier caliper and calculated using the following formula: tumor volume = (length x width^2^)/2. The mice were euthanized at 14 days after implantation, and the collected tumors were fixed with 10% formalin.

### Statistical Analysis

Statistical analyses were performed using R v4.4.1. For survival analysis, overall survival and progression-free survival curves were plotted according to the Kaplan-Meier method and compared using the log-rank test. For continuous variables, pairwise comparisons between groups were performed by a two-sided unpaired t-test. Categorical variables were compared using Fisher’s exact test. The differences were considered significant when the p-value was lower than 0.05. Data were represented as mean ± standard error of the mean (SEM).

### Data Availability

The RNA-seq datasets generated in this study have been deposited in the NCBI GEO with accession number GSE306447. The significant called peak information in the ChIP-seq data is available upon reasonable request.

## Results

### YAP/TEAD-driven transcriptional programs are hyperactivated in EACSCC

To clarify transcriptomic aberrations of EACSCC in an unbiased manner, we first performed RNA sequencing (RNA-seq) utilizing RNA extracted from 17 treatment-naïve primary EACSCC and 8 noncancerous ear skin tissue specimens from 17 patients with EACSCC (**Fig.1A**, Table S1). Principal component analysis (PCA) showed a clear separation between EACSCC and skin tissues, suggesting their distinct transcriptional profiles (**Fig. 1B** and **Fig. S1A**). We detected 3412 differentially expressed genes (DEGs), including 2052 upregulated- and 1370 downregulated-genes in EACSCC (**Fig. 1C**; absolute value of fold change > 2, FDR-adjusted p-value < 0.01). Gene ontology (GO) analysis of upregulated DEGs showed significant enrichment of genes involved in inflammatory responses, epithelial-mesenchymal transition (EMT) and cell cycle (**Fig. 1D**). Similarly, gene set enrichment analysis (GSEA) indicated that pathways involved in inflammation, EMT and cell cycle progression are positively enriched in primary EACSCC (**Fig. 1E**). Strikingly, previously established TEADi target signature in HNSCC (23), as well as YAP transcriptional target genes (24) were strongly enriched in EACSCC, suggesting the prominent role of YAP-driven transcriptional programs in EACSCC (**Fig. 1F**). In addition to upregulation of curated YAP/TEAD transcriptional target genes, several conventional or partial EMT (p-EMT) associated genes (33), as well as immune-evasive chemokines/cytokines, immune checkpoint molecules (*CD274* and *PDCD1LG2*) were overexpressed, whereas differentiation-related genes were downregulated (**Fig. 1G**). Consistently, activated transcription factors in EACSCC included TEAD1 and TEAD4 (**Fig. 1H**). Of note, survival analysis of these 17 patients indicated that TEADi target signature predicted poor survival of EACSCC patients, suggesting that YAP/TEAD transcriptional activity is correlated with malignant phenotypes of EACSCC (**Fig. 1I**). Together, transcriptome profiling suggested hyperactivation of the YAP/TEAD oncosignaling network in EACSCC.

**Figure 1.**
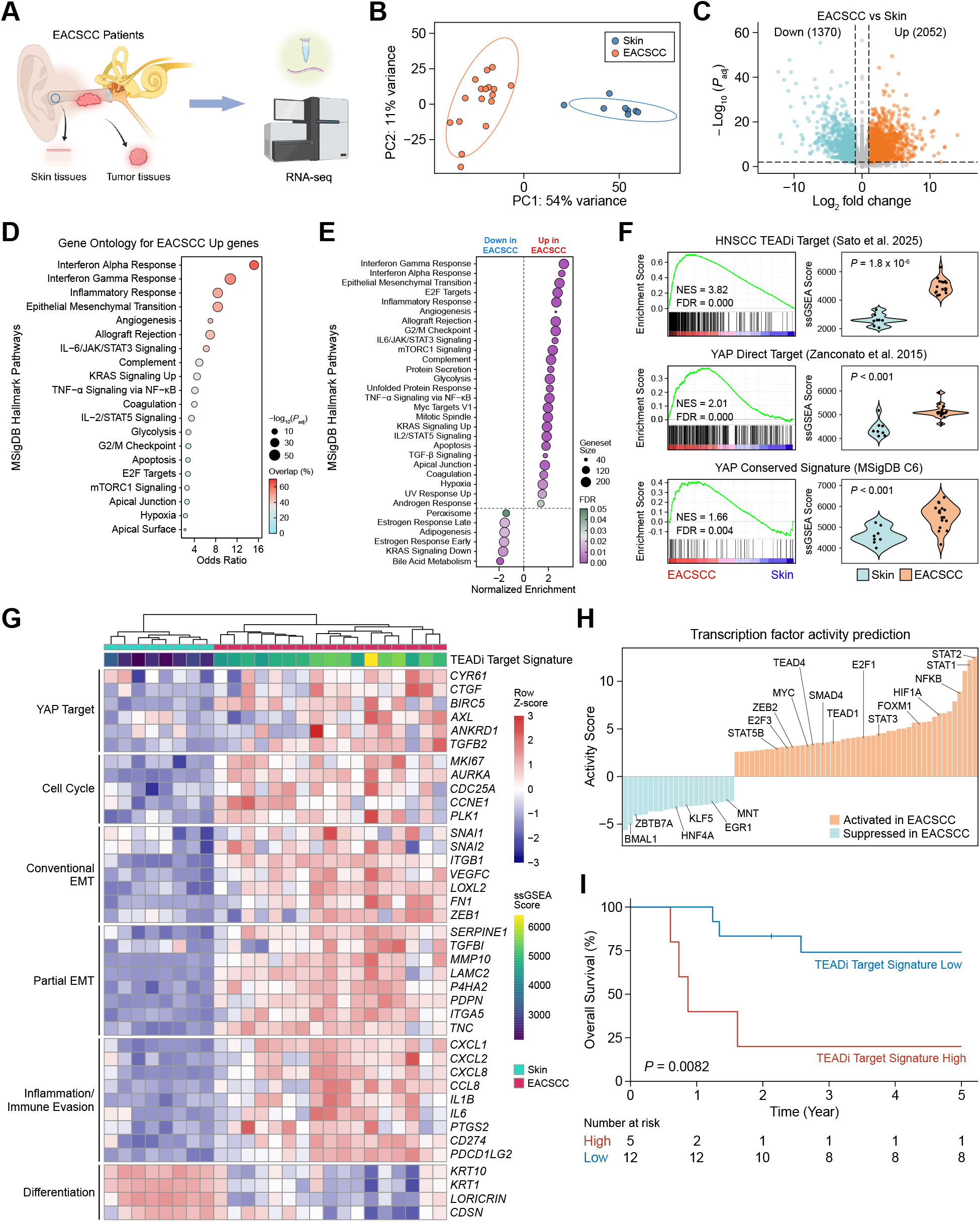
Transcriptomic analysis revealed hyperactivation of YAP signaling in EACSCC. **A**. Schematic representation for the experimental design of RNA-seq in EACSCC and noncancerous skin tissues. **B**. Principal component analysis for the RNA-seq data derived from EACSCC (n=17) and noncancerous skin tissues (n=8). **C**. Volcano plot representing differentially expressed genes (DEGs) between EACSCC and noncancerous skin tissues. **D**. Gene ontology (GO) analysis for the DEGs upregulated in EACSCC in Molecular Signature Database (MSigDB) Hallmark pathways. **E**. Gene set enrichment analysis (GSEA) representing gene sets positively or negatively enriched in EACSCC compared with noncancerous skin tissues in MSigDB Hallmark pathways. **F**. GSEA enrichment plots for the indicated YAP/TAZ-TEAD transcriptional target gene sets showing the enrichment between EACSCC and noncancerous skin tissues (left panels). Violin plots for single sample GSEA (ssGSEA) scores for each group are shown in right panels. **G**. The heatmap and hierarchical clustering for the selected genes in the indicated pathways. Gene expression levels were scaled among samples for each gene. The ssGSEA scores for the TEAD inhibitor target signature (Sato et al. 2025) are shown in the top of the heatmap. **H**. Waterfall plots showing the predicted transcription factor activities in EACSCC. **I**. Kaplan-Meier overall survival curves for the EACSCC patients with TEAD inhibitor target signature-high or low scores calculated by ssGSEA.

### YAP-driven epigenetic aberrations in EACSCC

Changes in chromatin accessibility and aberrant transcriptional machineries, including super enhancers (SEs), are one of the hallmarks of cancer (6,34), and YAP is a potential regulator for this reprogramming (13,35,36). Considering this point, we performed ChIP-seq utilizing an antibody for H3K27Ac, a histone mark for active enhancers and promoters, as well as YAP in clinical EACSCC tissues and noncancerous ear skin tissues to clarify upstream mechanisms that regulate aberrant transcriptional programs in EACSCC (**Fig. 2A**). We successfully performed H3K27Ac ChIP-seq in 5 EACSCC samples and 2 noncancerous ear skin samples, detecting 53400 peaks in EACSCC (range; 46038-64871), whereas 51311 in skin (range; 42967-59655) on average. PCA analysis of genome-wide H3K27Ac signal intensities showed a clear separation between EACSCC and skin, suggesting their distinct distribution of active transcription-regulatory elements (**Fig. 2B**). Based on these data, we predicted nucleosome-free regions (NFRs), accessible regulatory regions in enhancers and promoters, using a previously described bioinformatics algorithm. Motif enrichment analysis of NFRs showed significant enrichment of binding motifs for several TFs, such as AP-1 family (e.g., JUN and FOS), p63, TEAD, and Paired Like Homeodomains (PITX), which is reportedly involved in tissue regeneration and stemness (4,37,38) in EACSCC compared to skin, suggesting increased chromatin accessibility for these TF binding sites in this malignancy (**Fig. 2C**). We detected 1024 and 1021 SEs in EACSCC and skin on average, respectively (**Fig. S3A**). We next calculated overlapping SE regions between samples and defined tumor-specific SEs, which are exclusively observed in EACSCC tissues, to explore their possible biological roles in EACSCC (see Method) (**Fig. 2D**). GO analysis for the putative transcriptional target genes of EACSCC-specific SEs indicated enrichment of oncogenic processes such as EMT, cell cycle, hypoxia and inflammatory response, suggesting that upregulated genes in EACSCC tissues that we identified above (**Fig. 1**) could be regulated in part through the SEs (**Fig. 2E**). Subsequently, we sought to clarify YAP-driven transcriptional aberrations in EACSCC utilizing YAP ChIP-seq, and integrating with H3K27Ac ChIP-seq data. We were able to perform YAP ChIP-seq on 2 EACSCC tissues and paired 2 noncancerous skin tissues. Clustering analysis of YAP signals around YAP peaks indicated EACSCC-specific binding patterns of YAP (Cluster 2; **Fig. 2F**). As expected, motif analysis for 200bp regions around YAP1 peak summits in this cluster showed significant enrichment of DNA binding motifs for TEAD and AP-1, indicating that these TFs are predominantly forming complexes with YAP as previously shown in cultured cells (24,39) (**Fig. 2G**). Notably, GO analysis of predicted YAP target genes showed strong enrichment of EGFR signaling, proliferation, and differentiation in EACSCC, which were not observed in skin (**Fig. 2H**). Unexpectedly, we observed significant YAP binding peaks with SE formation around the coding region for *EGFR* exclusively in EACSCC, suggesting that YAP/TEAD may directly regulate *EGFR* transcription through SEs (**Fig. 2I**). In addition, YAP binding peaks were observed in *EREG* (Epiregulin) and *AREG* (Amphiregulin), both of which are ligands for EGFR, as well as EMT-promoting TF *SNAI2* and immune evasive chemokine *CXCL8* (**Figs. S3B-D**). Overall, these results indicated YAP could drive oncogenic transcriptional programs thorough SEs in EACSCC.

**Figure 2.**
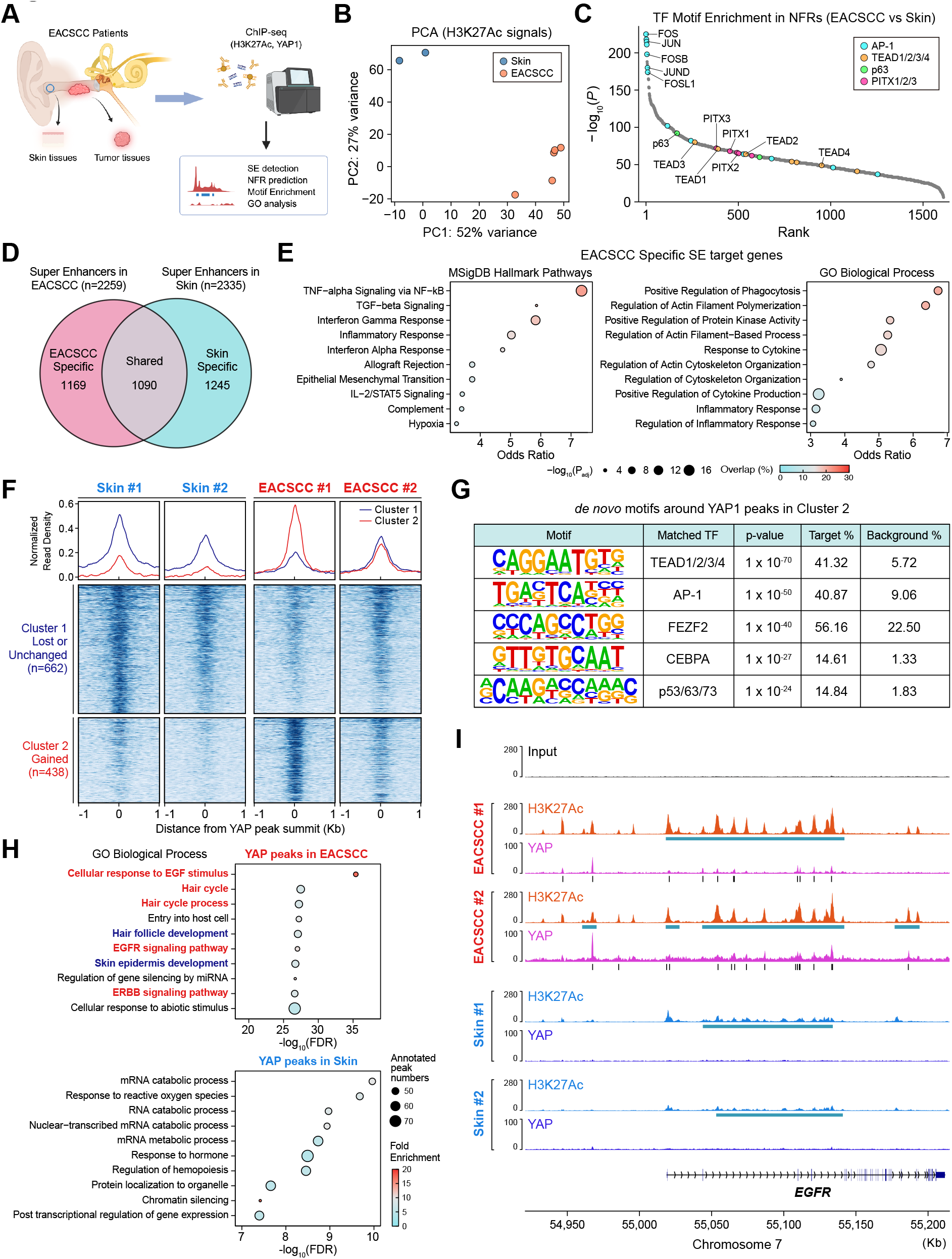
YAP-driven epigenetic reprogramming in EACSCC. **A**. Schematic representation for the experimental design of ChIP-seq in EACSCC and noncancerous skin tissues. **B**. Principal component analysis for the genome-wide H3K27Ac ChIP-seq signals derived from EACSCC (n=5) and noncancerous skin tissues (n=2). **C**. Enrichment of transcription factor (TF) binding motifs in putative nucleosome free regions (NFRs) in EACSCC compared with noncancerous skin tissues. Selected TF motifs are highlighted. **D**. Venn diagrams for the super enhancers (SEs) in EACSCC and noncancerous skin tissues. **E**. GO analysis for EACSCC-specific SE target genes in MSigDB Hallmark Pathways (left) and GO Biological Process (right). **F**. Read density plots and heatmaps for YAP ChIP-seq signals in ±1□kb windows centered at YAP peak summits in EACSCC and noncancerous skin tissues. K-means clustering (k=2) was performed to detect lost/unchanged (cluster 1, n=662) and gained (cluster 2, n=432) YAP signals in EACSCC. **G**. *de novo* TF binding motif discovery analysis in 200bp regions around YAP ChIP-seq peaks in the cluster 2. **H**. GO analysis for the predicted transcriptional target genes of YAP in EACSCC and noncancerous skin tissues. Pathways involved in EGFR signaling and proliferation are highlighted in red, whereas pathways involved in differentiation are highlighted in blue. **I**. Genome browser snapshot representing H3K27Ac and YAP occupancy at the *EGFR* gene locus in EACSCC and noncancerous skin tissues. Detected SE regions and YAP peak summits are shown in blue and black bars, respectively.

### PITX2, a potential alternative partner TF of YAP under TEAD-inhibited condition, regulates oncogenic transcription in EACSCC

Our sequencing data provided a rationale for hyperactivation of YAP in EACSCC, and suggested that EACSCC could be dependent on YAP-driven transcriptional programs. Based on the results of YAP ChIP-seq in our clinical EACSCC samples, YAP predominantly interacts with TEAD as previously reported *in vitro* and *in vivo* (40), suggesting that pharmacological inhibition of YAP/TEAD interaction may serve as a potential therapeutic approach of EACSCC. We sought to explore this question utilizing VT104, a small molecule TEAD auto-palmitoylation inhibitor which blocks YAP-TEAD interaction (41), in EACSCC-derived cells SCEAC-ono2 (referred to as ONO2 hereafter) previously established from primary EACSCC (31). As expected, VT104 inhibited proliferation and clonogenicity of ONO2 (**Figs. 3A and 3B**), and several YAP/TEAD target genes were downregulated in qPCR upon VT104 treatment (**Fig. 3C**), indicating the dependencies on YAP/TEAD signaling in EACSCC cells. However, whether alternative TFs could bind to YAP and be involved in oncogenic transcription under certain conditions is still controversial. Motif analysis in NFRs indicated that binding sites for several TFs are accessible in EACSCC but not in skin (**Fig. 2C**), suggesting the possibility that YAP may interact with TFs other than TEAD. One of the most significantly enriched TF binding motifs in accessible regions on enhancers included PITX2 transcription factor, which is reportedly involved in heart muscle regeneration cooperatively with Yap1 in mice (37), as well as wound healing (4), oncogenic transcription and stemness in cancer (38). Thus, we hypothesized that PITX2 could be an alternative partner TF of YAP and performed further analyses. Strikingly, YAP bound to PITX2 upon VT104 treatment, while dissociation between YAP and TEAD1 was observed in co-immunoprecipitation assays (**Fig. 3D**). Interestingly, binding between YAP and PITX2 was also observed in HSC4 HNSCC cells without TEAD inhibition, possibly indicating a more predominant role of this TF in HNSCC (**Fig. S4A**). Knockdown of PITX2 significantly reduced proliferation and clonogenicity, as well as migration in ONO2 and representative HNSCC cells FaDu, HSC4, and SCC9 (**Figs. 3E-F and S4C-E**). To clarify the oncogenic function of PITX2 in EACSCC, we next performed RNA-seq in ONO2 cells stably overexpressing PITX2 and control cells. Notably, PITX2 induced expression of genes involved in partial- or conventional-EMT (*VIM, LAMC2, LAMA3, MMP10* and *FN1*), stemness (*CD44*), and cell cycle progression (*MYC, CCNE1, CCND1* and *CDC25A*). Unexpectedly, significantly upregulated genes included well-characterized YAP/TEAD target genes (*CYR61, CTGF*, and *ANKRD1*), suggesting that PITX2 may partially rescue YAP/TEAD-driven transcription (**Fig. 3G**; absolute value of fold change > 1.5, FDR-adjusted p-value < 0.01). GO analysis and GSEA indicated significant enrichment of genesets involved in cell cycle progression and EMT (**Figs. 3H and 3I**). Consistently, we observed upregulation of CD44, as well as EMT markers CDH2 and SNAI1 upon PITX2 overexpression by western blotting (**Fig. 3J**). Of note, PITX2 overexpression significantly promoted proliferation, migration, and spheroid formation of ONO2 *in vitro* (**Figs. 3K-3M**), as well as tumor growth *in vivo* (**Figs. 3N-3P**). These results suggested the oncogenic transcriptional ability of PITX2 in EACSCC.

**Figure 3.**
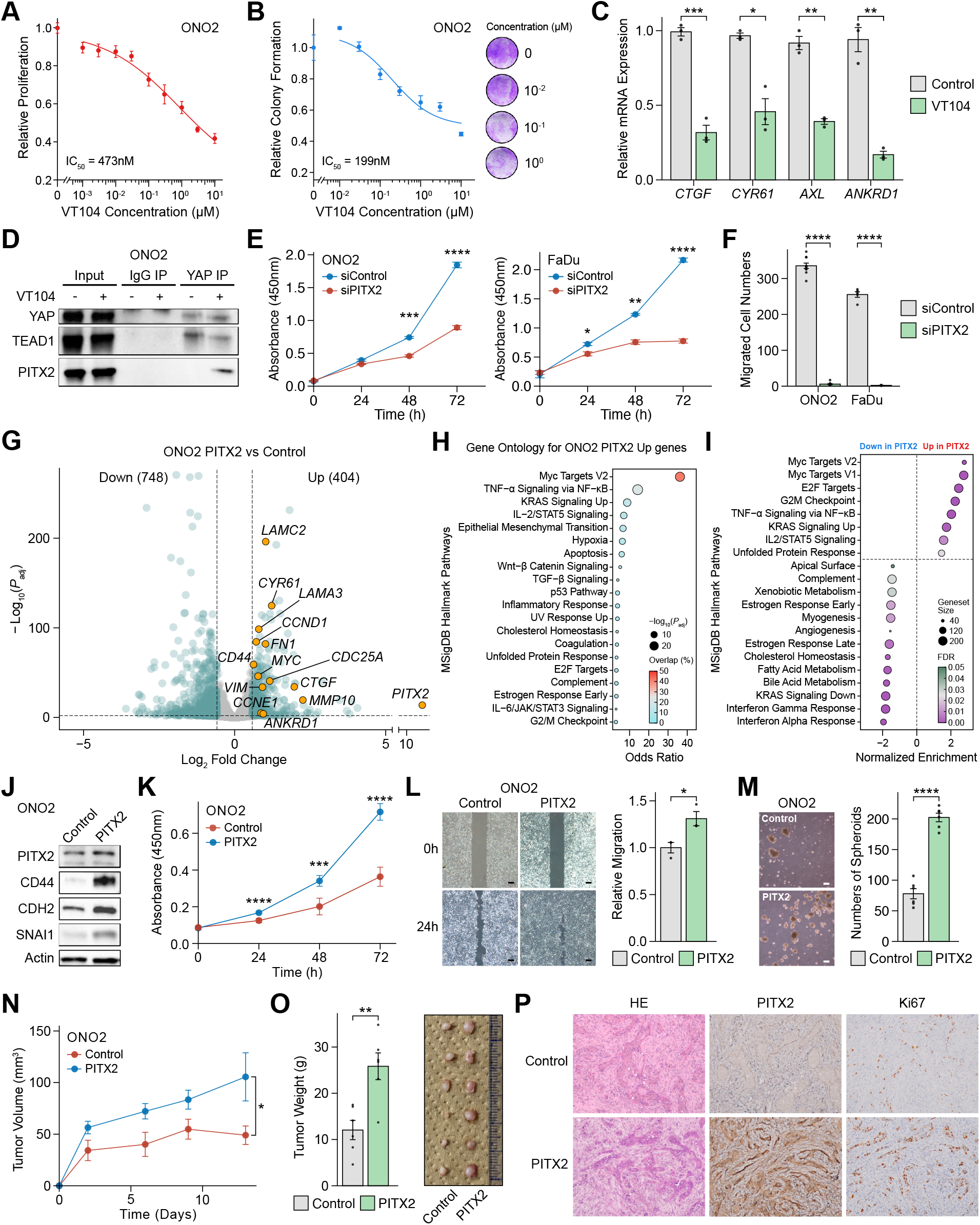
PITX2 acts as potential partner transcription factor of YAP in EACSCC. **A and B**. Dose response curves of VT104 in SCEACono2 (ONO2) cells examined in proliferation assay (**A**) and clonogenic assay (**B**). **C**. qPCR for YAP/TAZ-TEAD target genes upon treatment with VT104 (1*µ*M) for 24 hours in ONO2 cells *in vitro* (n=3). **D**. YAP co-immunoprecipitation experiment in ONO2 cells treated with VT104 (1*µ*M) for 24 hours. **E**. Proliferation assays in ONO2 (left) and FaDu (right) upon siRNA-mediated knockdown against PITX2 (n=3). F. Trans-well migration assays in ONO2 and FaDu upon siRNA-mediated knockdown against PITX2 (n=5). **G**. Volcano plot representing DEGs between PITX2-overexpressed and control ONO2 cells. Selected genes are highlighted as yellow dots.Genes with adjusted P-value < 0.01 and absolute value of fold change > 1.5 were considered as significant. **H**. GO analysis for the DEGs upregulated in PITX2-overexpressed ONO2 cells compared with control cells in MSigDB Hallmark pathways. **I**. GSEA representing gene sets positively or negatively enriched in PITX2-overexpressed ONO2 cells compared with control cells in MSigDB Hallmark pathways. **J**. Western blots for the indicated proteins in PITX2-overexpressed and control ONO2 cells. **K**. Proliferation assays in PITX2-overexpressed and control ONO2 cells (n=5). **L**. Wound healing assays in PITX2-overexpressed and control ONO2 cells (n=3). **M**. Spheroid formation assays in PITX2-overexpressed and control ONO2 cells (n=6). **N and O**. Growth curves for PITX2-overexpressed or control ONO2 xenograft tumors (**N**) and barplots for tumor weight at endpoint (**O**) (n=6). The image of tumors is also shown. **P**. Representative images for HE staining as well as immunohistochemical staining for PITX2 and Ki67 in the xenograft tumors. Data represent mean ± SEM. ^*^*P* < 0.05, ^**^*P* < 0.01, ^***^*P* < 0.001, ^****^*P* < 0.0001.

### YAP and PITX2 predict poor prognosis of EACSCC patients

Finally, we examined the expression of YAP protein in treatment-naïve biopsy or surgically resected primary tumor tissue samples from 74 EACSCC patients by immunohistochemistry (IHC). Overexpression of YAP was observed in the nucleus of tumor cells in 40 out of 74 specimens examined (54%), suggesting that YAP is in its functionally active state (**Fig. 4A**). EACSCC patients with high expression levels of YAP showed significantly poor overall and progression-free survival (**Figs. 4B and 4C**). We also evaluated PITX2 expression in EACSCC tissues (N=79), and found that PITX2 was highly expressed in the nucleus of EACSCC cells in 22 out of 79 specimens (28%) (**Fig. 4D**). Of note, high PITX2 expression was positively correlated with poor overall and progression-free survival, similar to the result of YAP IHC (**Figs. 4E and 4F**). In addition, YAP and PITX2 expression levels were positively correlated in the overlapping cases (N=73; Fisher’s exact test, *P*□=□0.022) (**Fig. S5**). These clinical data may support our sequencing and experimental findings, indicating that both YAP and PITX2 are prognostic factors for EACSCC.

**Figure 4.**
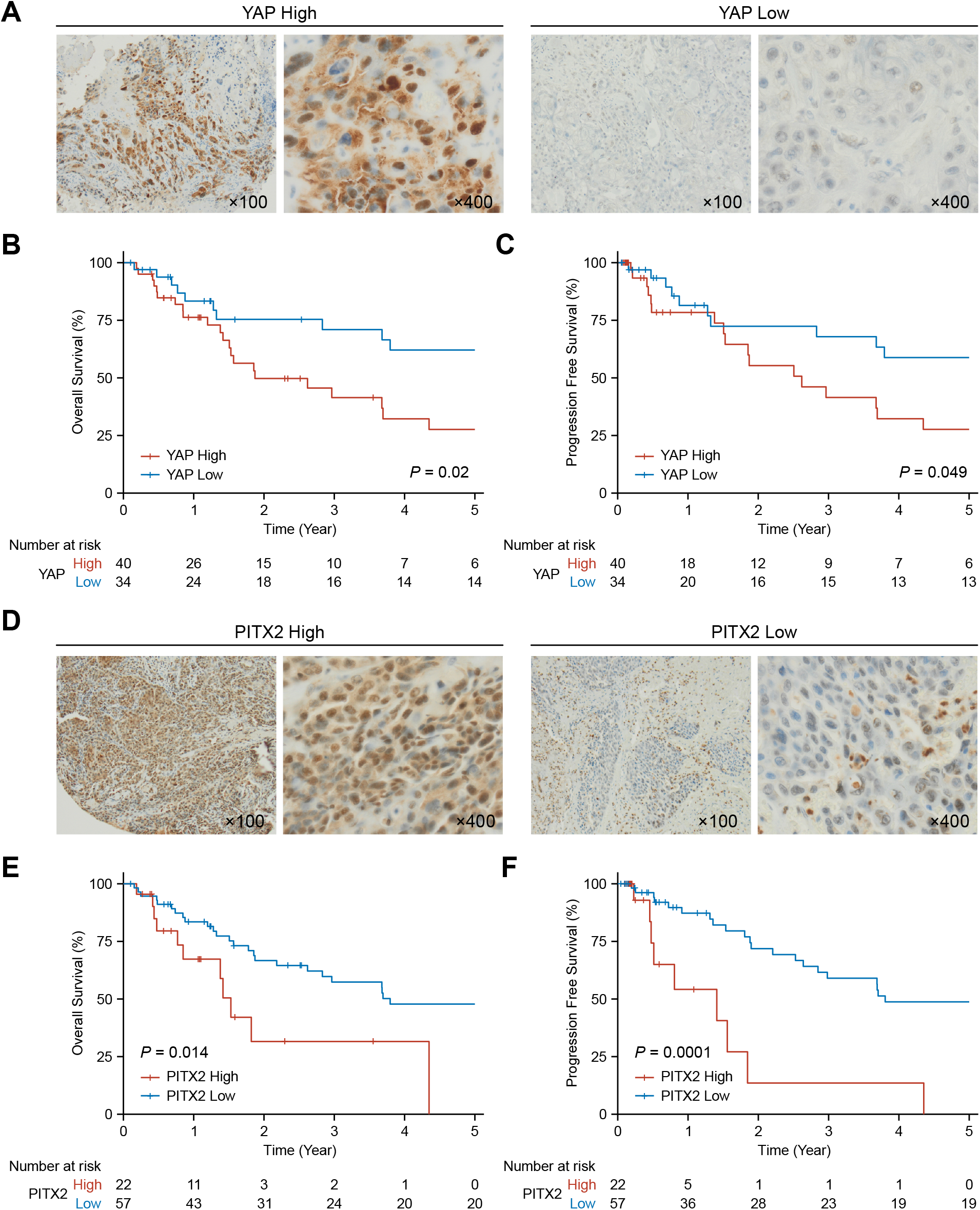
YAP and PITX2 predict poor prognosis of EACSCC patients. **A**. Representative images of immunohistochemical staining for YAP in EACSCC tissues. **B and C**. Kaplan-Meier curves for overall survival (**B**) and progression free survival of EACSCC patients classified according to the YAP expression levels. **D**. Representative images of immunohistochemical staining for PITX2 in EACSCC tissues. **E and F**. Kaplan-Meier curves for overall survival (**B**) and progression free survival of EACSCC patients classified according to the PITX2 expression levels. P-values were calculated using log-rank test.

## Discussion

The rarity of EACSCC has prevented the development of evidence-based therapeutic strategies as well as a comprehensive understanding of its biological and molecular backgrounds. One of the unique aspects of EACSCC is that this malignancy could be induced by chronic tissue damage and inflammation. This may be supported by our previous genomic study of EACSCC, in which tobacco- or UV-induced mutational signatures were not evident; instead, we identified predominant APOBEC signatures, which suggested its link to chronic inflammation (17), similar to aggressive cutaneous SCC in recessive dystrophic epidermolysis bullosa driven by inflammatory microenvironment (42). Considering that aberrant and continuous hyperactivation of regenerative transcriptional programs in response to chronic tissue damage may fuel oncogenesis (6,43), EACSCC could be addicted to this oncogenic transcription, and thus, this could be a therapeutic vulnerability that can be exploited for therapeutic purposes as well. However, unlike other types of malignancies in which transcriptomic and pathway level alterations are well described in large-scale studies, the core transcriptional programs activated in EACSCC have not yet been clarified.

In this study, we provided evidence of hyperactivation of YAP-driven transcriptional programs in EACSCC. Indeed, RNA-seq analysis indicated that curated YAP/TEAD target genesets were significantly enriched in EACSCC (23,24), which was supported by ChIP-seq and IHC analyses of clinical samples. Notably, YAP/TEAD-driven transcriptional programs, as well as YAP nuclear expression, were correlated with poor prognosis of EACSCC patients, suggesting that YAP hyperactivation may drive the malignant phenotype of EACSCC.

To date, no study has achieved YAP ChIP-seq in human clinical tissue samples. Our YAP ChIP-seq data may provide a better rationale for hyperactivation of YAP/TEAD in human malignancies and suggest its downstream pathways, including EMT, chemokine production, and EGFR signaling. Specifically, SE formation and YAP binding in *EGFR*, as well as its ligands *EREG* and *AREG* loci, indicated that YAP/TEAD may directly control the transcription of these targets through SE. These findings may support the potential positive-feedback loop between YAP/TEAD and EGFR signaling, as recent studies have shown that YAP/TEAD promotes transcription of EGFR ligand *NRG1* (13) and that EGFR activates YAP/TEAD through phosphorylation of Hippo component (44). Moreover, H3K27Ac ChIP-seq indicated genome-wide SE formation and gained accessibility for oncogenic TF binding sites in EACSCC. Previous studies demonstrated that YAP recruits bromodomain-containing protein 4 (BRD4), an epigenetic co-factor essential for SE formation to enhancer and maintains active chromatin state (35,36,45), suggesting that YAP could be a master regulator for SE-mediated oncogenic transcription. In this context, the co-occurrence of SE and YAP bindings exclusively observed in EACSCC may indicate the crucial role of YAP-driven epigenetic reprogramming and transcriptional addiction to Hippo-YAP/TEAD oncogenic signaling network in EACSCC.

Unexpectedly, we found that YAP bound to PITX2, a TF involved in stemness and tissue regeneration (37,38), under pharmacological YAP/TEAD blockade. Generally, TEAD is considered the most important YAP-binding TF, whose interacting mechanisms are well documented by a series of biological studies (9,40), which ultimately resulted in the discovery and development of small molecule TEAD inhibitors (7,23,41,46). However, the mechanistic consequence of YAP/TEAD inhibition is not yet fully uncovered, which may be context-dependent and cell-type specific (47). Our Recent studies demonstrated that YAP/TEAD inhibition induced terminal differentiation of HNSCC and skin epithelial cells, which could be driven by other TFs such as KLF4, suggesting alternative transcriptional machineries other than YAP/TEAD (23,48). Although the exact mechanism of YAP/PITX2 interaction still needs to be further elucidated, our findings suggest that PITX2 may rescue YAP/TEAD-driven oncogenic transcription, including proliferation, EMT, and stemness. Importantly, the function of PITX TFs has been mainly described in the context of tissue regeneration and wound healing (4,37). Of interest, both PITX1 and PITX2 are expressed in oral mucosa and involved in rapid wound healing, whereas these TFs are not expressed in normal skin, even in the regeneration process (4). In this regard, our experimental data, as well as the overexpression of YAP and PITX2 in EACSCC tissues, suggest a transcriptionally reprogrammed state of EACSCC and its unique oncogenic transcriptional networks, which may propose a potential therapeutic strategy co-targeting YAP/TEAD and YAP/PITX2 signaling axes.

Of note, IHC analysis of PITX2 demonstrated that nuclear expression of PITX2 protein significantly correlated with poor overall and progression-free survival, suggesting its contribution to aggressive phenotypes of EACSCC and potential as a prognostic biomarker of EACSCC patients, which was further supported by our functional experiments on PITX2 in EACSCC and HNSCC cells. Interestingly, *PITX2* mRNA expression as well as hypomethylation of *PITX2* promoter were reportedly correlated with poor prognosis of HNSCC patients (49,50). Similarly, PITX2 promoted stemness and proliferation of esophageal SCC (ESCC) cells and predicted poor prognosis of ESCC patients (38), all of which may suggest its potential oncogenic role in squamous cancer.

In summary, we describe the transcriptomic and epigenetic aberrations of EACSCC and provide evidence that hyperactivated YAP mediates transcriptional programs that drive the malignant phenotypes in EACSCC. Our findings may contribute to a better understanding of YAP-driven epigenetic reprogramming not only in EACSCC but also in human squamous malignancies, as well as provide a mechanistic rationale for the future clinical development of novel therapeutic strategies, such as those targeting the Hippo-YAP/TEAD signaling network, for EACSCC patients.

## Acknowledgments

K.S. was supported by Japan Society for the Promotion of Science (JSPS) Grant 20K18300, SGH Foundation Cancer Research Grant, Shinnihon Foundation of Advanced Medical Treatment Research Grant, Soda Toyoji Memorial Foundation Research Grant, Takeda Science Foundation Research Grant and Takeda Science Foundation Research Fellowship. N.K. was supported by JSPS Grant 22K09745. T. Nakagawa was supported by JSPS Grant 22H03236.

This work used the supercomputing resources provided by the Human Genome Center, Institute of Medical Science, University of Tokyo (http://sc.hgc.jp/shirokane.html).

Conception and design: K.S.

Acquisition of Data: K.S., N.K., M.O., T.H., T. Nakano, K.K. and K.T.

Data Analysis: K.S., N.K., M.O., T.H., T. Nakano and S.I.

Writing the original manuscript: K.S.

Writing and Review of the manuscript: K.S., N.K., J.S.G., K.M., M.M. and T.

Nakagawa Supervision: K.S. and N.K. and T. Nakagawa

All cartoon renderings were created with the BioRender online platform (BioRender.com)

