## Supplementary material for "Aberrant Hippo-YAP/TEAD signaling drives malignant transcriptional reprogramming in external auditory canal squamous cell carcinoma": Figure S1

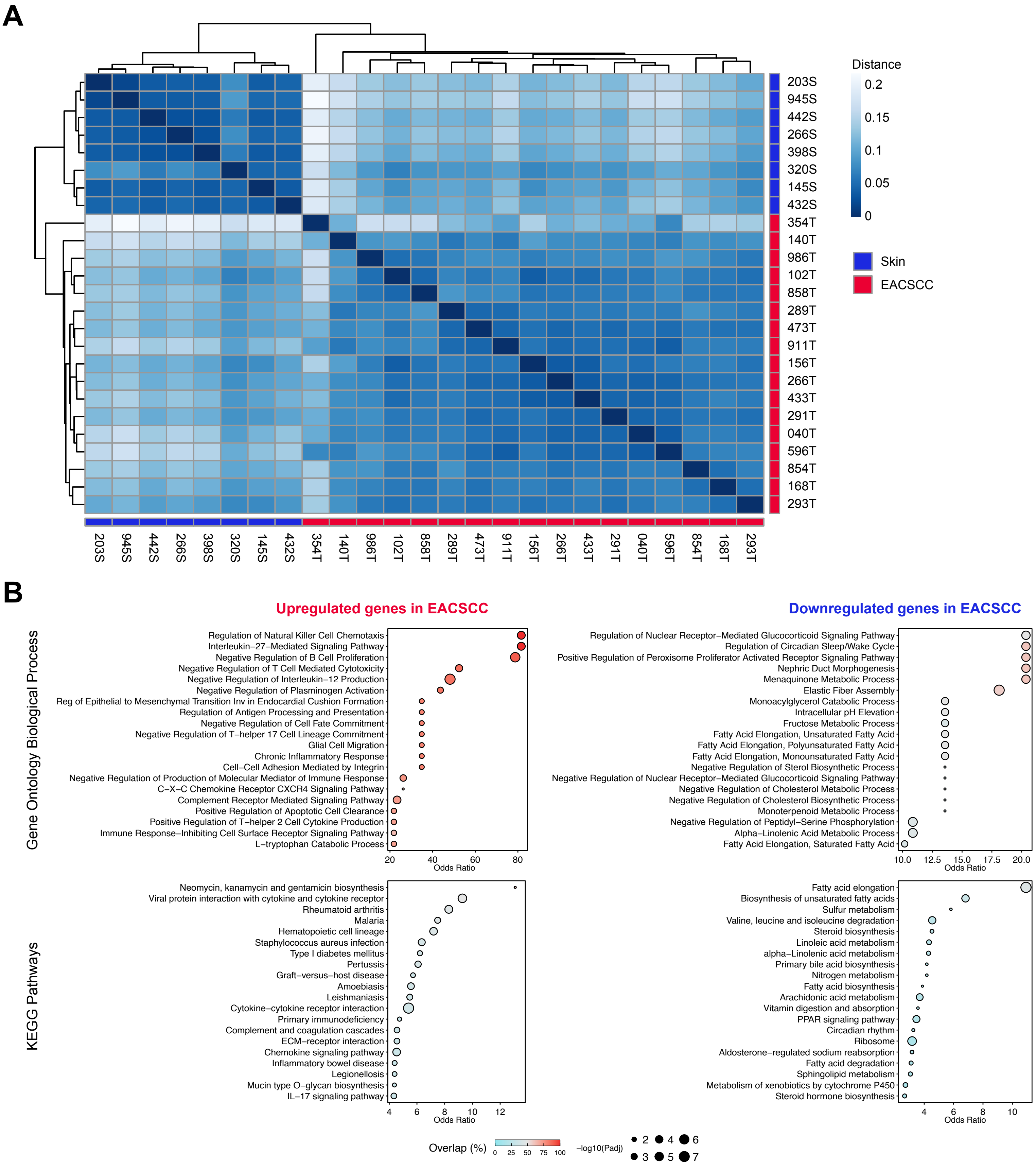


**Figure S1. Transcriptomic landscapes of EACSCC is distinct from noncancerous ear skin tissues.** **(A)** Transcriptomic distances of EACSCC (n=17) and noncancerous ear skin tissues (n=8). **(B)** Gene ontology analysis in Gene Ontology Biological Process (upper panels) and KEGG Pathways (lower panels) for the upregulated (left) and downregulated (right) genes in EACSCC compared with noncancerous ear skin tissues.
