## Supplementary material for "Aberrant Hippo-YAP/TEAD signaling drives malignant transcriptional reprogramming in external auditory canal squamous cell carcinoma": Figure S2

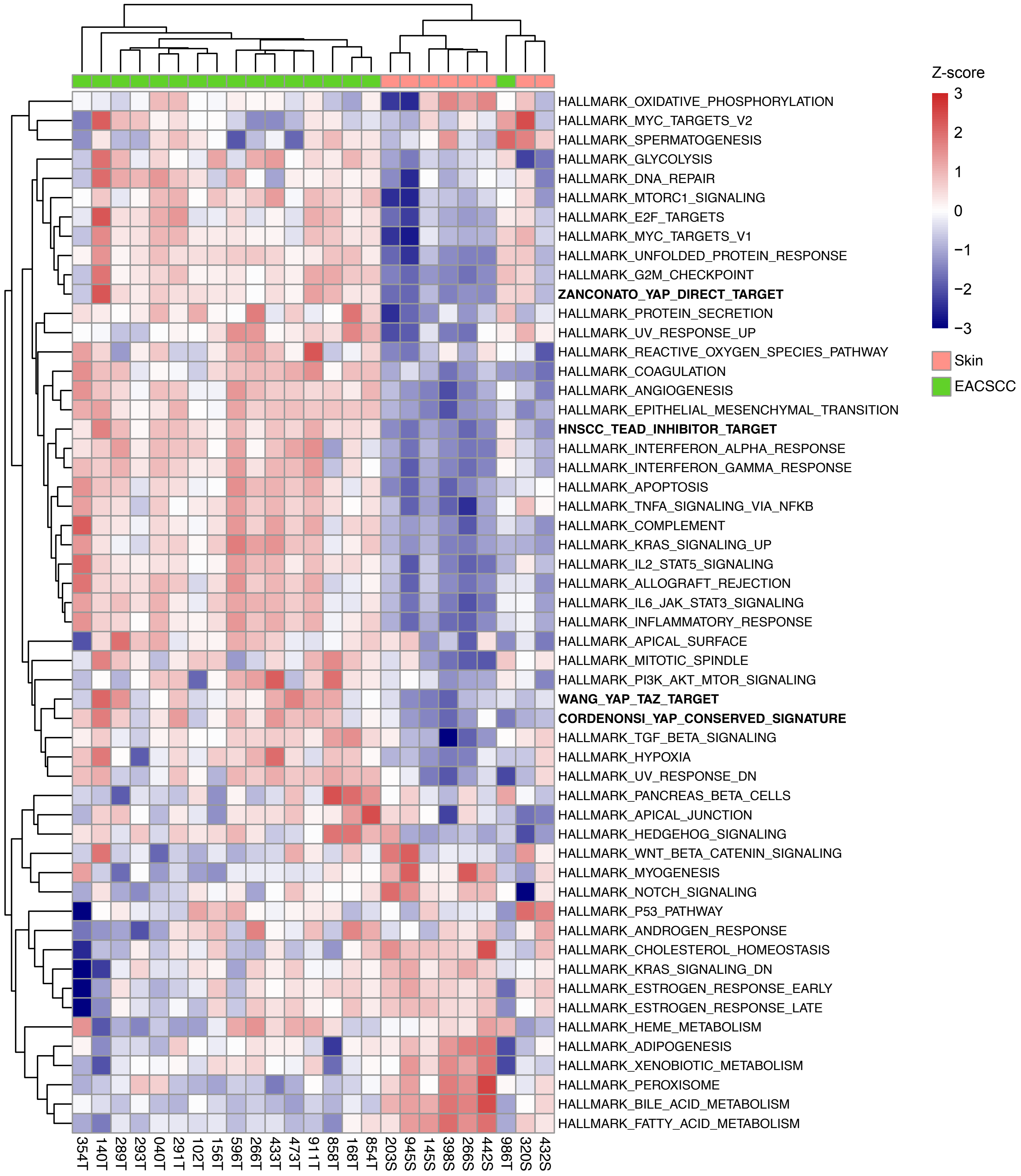


**Figure S2. Pathway-level transcriptomic aberrations of EACSCC.**

The Heatmap for the scores from single sample Gene Set Enrichment Analysis in EACSCC and noncancerous ear skin tissues. Molecular Signature Database Hallmark genesets and YAP/TAZ-TEAD target genes were included for the analysis.
