## Supplementary material for "Aberrant Hippo-YAP/TEAD signaling drives malignant transcriptional reprogramming in external auditory canal squamous cell carcinoma": Figure S3

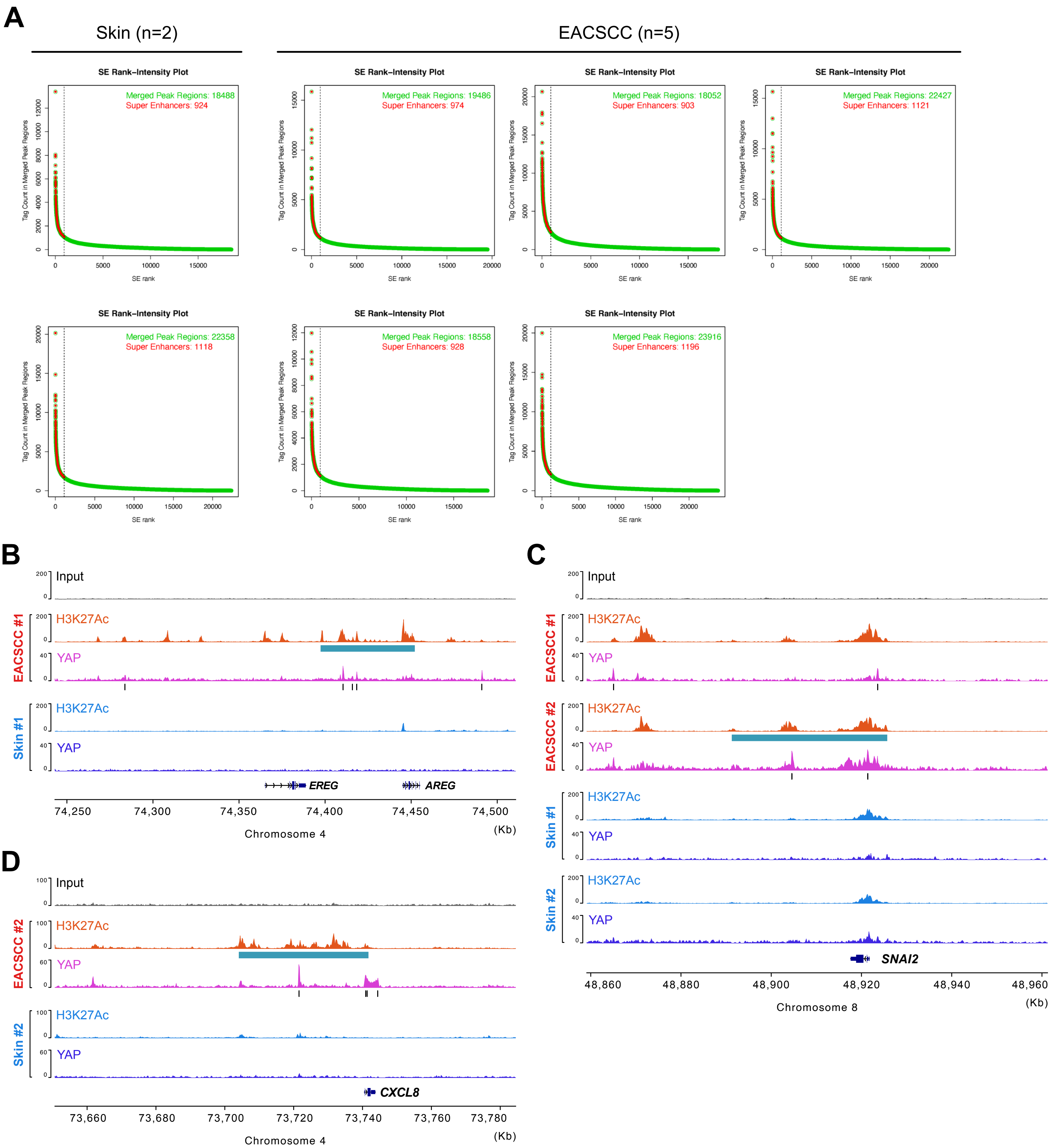


**Figure S3. Super Enhancer (SE) formation and YAP binding in EACSCC.**
**(A)** Enhancer regions (green) and SE regions (red) ranked by H3K27Ac signals at enhancers obtained by ROSE in noncancerous ear skin tissues (n=2) and EACSCC (n=5). **(B-D)** Genome browser snapshots for H3K27Ac and YAP occupancy at *EREG* and *AREG* (**B**), *SNAI2* (**C**) and *CXCL8* (**D**) coding regions. Blue and black bars represent SEs and YAP peaks, respectively.
