## Supplementary material for "Aberrant Hippo-YAP/TEAD signaling drives malignant transcriptional reprogramming in external auditory canal squamous cell carcinoma": Figure S4

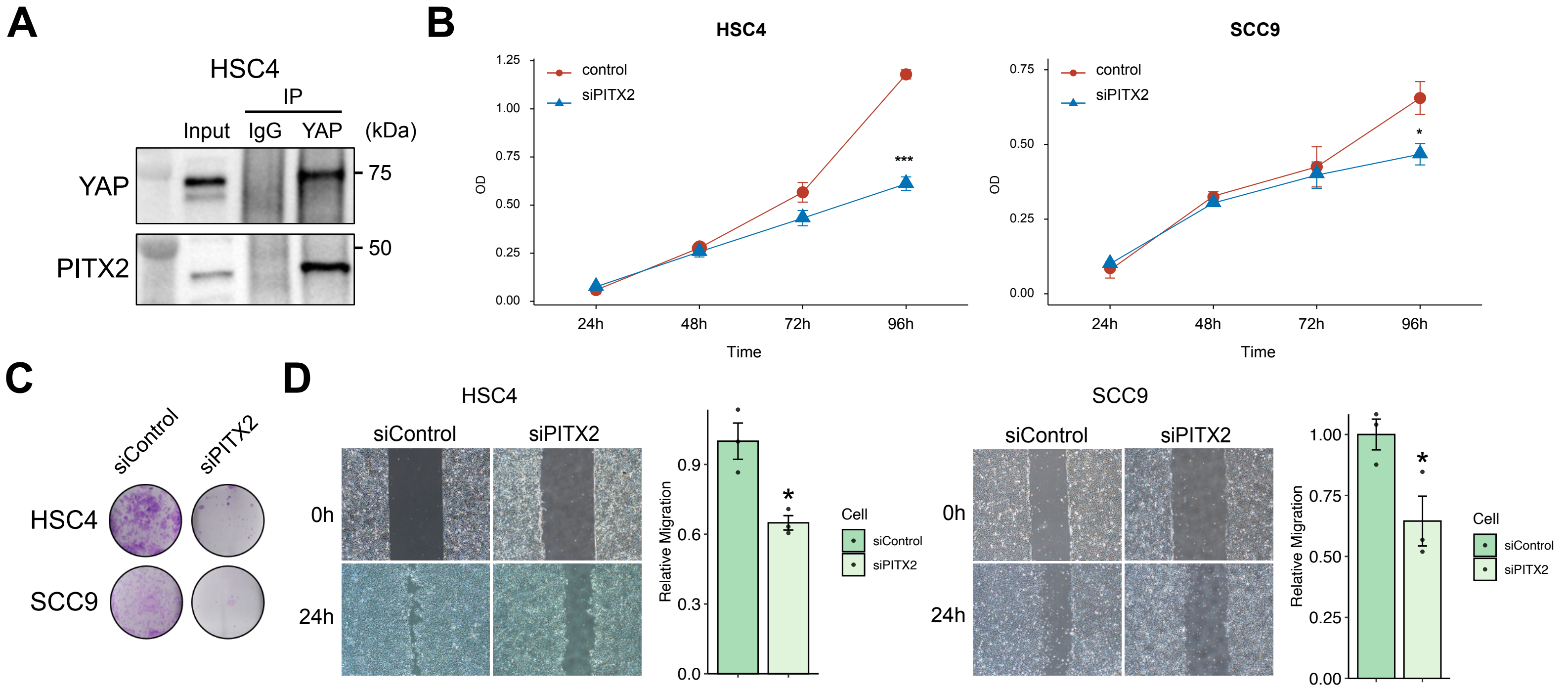


**Figure S4. Knockdown of PITX2 suppresses proliferation and migration of SCC cells *in vitro*.**
**(A)** Co-immunoprecipitation utilizing YAP antibody in HSC4 HNSCC cells. Western blot for YAP and PITX2 are shown. **(B)** Proliferation assays in HSC4 and SCC9 cells treated with siPITX2 and control siRNA. n=3 for each group. **(C)** Clonogenic assays in HSC4 and SCC9 cells treated with siPITX2 and control siRNA. **(D)** Migration assays in HSC4 and SCC9 cells treated with siPITX2 and control siRNA. n=3 for each group. *$P$ < 0.05, ***$P$ < 0.001.
