## Supplementary material for "Aberrant Hippo-YAP/TEAD signaling drives malignant transcriptional reprogramming in external auditory canal squamous cell carcinoma": Figure S5

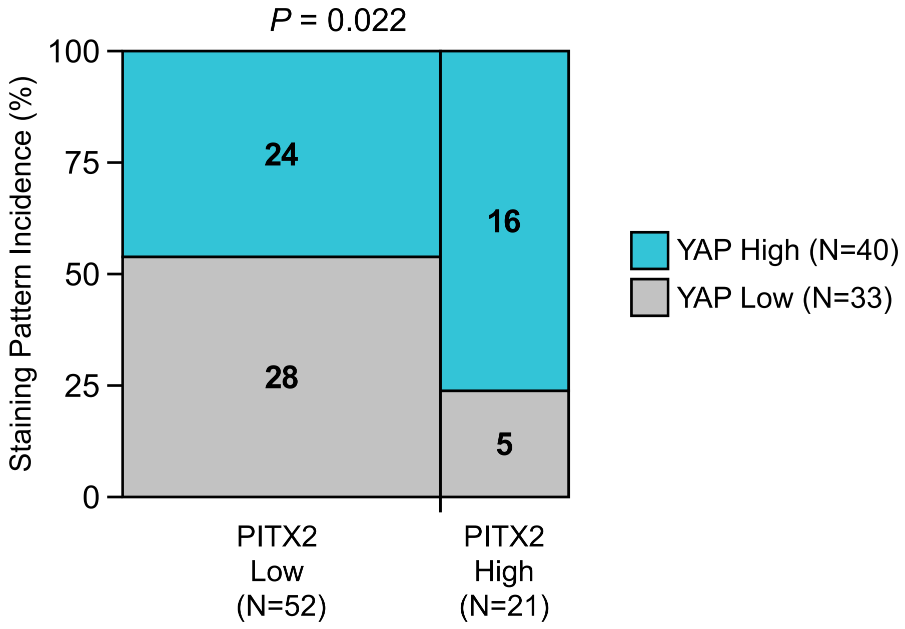


**Figure S5. YAP1 and PITX2 expression levels are positively correlated in EACSCC tissues.** Mosaic plot summarizing YAP and PITX2 expression in EACSCC tissues for the indicated number of EACSCC patients (N = 73). The p-value for the association between the parameters was calculated via Fisher's exact test.
